# Genomic regions exhibiting divergent methylation patterns covary with loci associated with mate choice traits in a stick insect

**DOI:** 10.1101/2025.07.30.667717

**Authors:** Clarissa F. de Carvalho, Nicholas P. Planidin, Jon Slate, Rüdiger Riesch, Zachariah Gompert, Patrik Nosil

## Abstract

Speciation involves the development of reproductive isolation between diverging populations. A potential key driver of reproductive isolation is mate choice, a behavioural mechanism that can limit gene flow based on divergence in signal traits. While the genetic basis of mating signal traits has been extensively studied, the contribution of epigenetic modifications to their variation remains underexplored, leaving the role of DNA methylation in mate choice unclear. Here, we focus on epigenetic variation and cuticular hydrocarbons (CHCs), the latter being chemical traits used for mate choice in insects. Specifically, we investigate the association between DNA methylation and regions associated with CHC variation in *Timema cristinae* stick insects. We integrate analyses of differentially methylated regions (DMRs) between individuals from different host-plant ecotypes with genomic sequencing and phenotypic data on CHCs. We find that DMRs are significantly enriched in genetic loci associated with CHCs, suggesting a non-random relationship between DNA methylation and loci associated with these signal traits. While further work is required to clarify causality, our results highlight the potential for epigenetic marks to be associated with traits involved in mate choice. Future studies should thus aim to establish causal links between DNA methylation and signal trait variation, which would clarify the contribution of methylation to mate choice, prezygotic isolation, and ultimately, speciation.

## Introduction

The origin of species involves the accumulation of phenotypic and genetic differences alongside the development of barriers that create reproductive isolation [1–3]. One major form of reproductive isolation is prezygotic isolation – barriers that act before fertilization, including temporal isolation, habitat isolation, mechanical isolation, sexual mating isolation, and gametic isolation [4]. A common type of prezygotic isolation is sexual isolation through mate choice, where individuals discriminate mates based on traits such as colouration, pheromones, songs, or other behaviours [4–9]. Mate choice arises from interactions between two types of traits: mating preferences of choosing individuals, and the signal traits of potential mates (mate choice traits, hereafter) [10,11]. This way, mate choice can limit gene flow when mating preferences differ between groups, contributing to the divergence of signal traits and sexual isolation. Understanding mate choice can therefore offer key insights into the mechanisms of prezygotic isolation and the evolution of reproductive barriers [12– 15].

Because mate choice directly affects reproductive success, it is a major driver of sexual selection. At the same time, both mate preferences and sexual signals can be sensitive to environmental and physiological conditions, causing context- and condition-dependent variation in trait expression [16,17]. As a result, the expression of signal traits and the preferences for such traits may vary in space and time [18,19]. Such phenotypic plasticity of sexual traits might even reflect the individual’s physiological condition and potentially inform potential mates about their quality [20–23], although not necessarily always the case [24,25]. A classic example of condition-dependent sexual signalling is the expression of colouration through dietary intake of carotenoids. Vibrant colours associated with higher carotenoid intake tend to be generally preferred – as observed in several species, such as the house finch (*Haemorhous mexicanus*) [26], guppies (*Poecilia reticulata*) [27], and sticklebacks (*Gasterosteus aculeatus*) [28].

Mate choice traits are thus shaped by both ecological factors and genetics [16,29]. In this context, numerous studies have examined the molecular basis of mate choice traits, an important step to understanding their evolution. A strong emphasis is often given to the genetic architecture of signal traits. Colouration has received particular attention across taxa, including plumage colour in capuchino seedeaters (*Sporophila* spp.) [30] and barn swallows (*Hirundo rustica*) [31], wing pattern in *Heliconius* butterflies [32,33], and colour traits in fish such as cichlids [34] and guppies [35]. Other types of traits have also received attention, such as songs in *Laupala* crickets [36] and *Pogoniulus* tinkerbirds [37], morphological features like swordtail extensions in *Xiphophorus* fishes [38] and body shape in sticklebacks (*Gasterosteus aculeatus*) [39], and chemical cues such as cuticular hydrocarbons in insects [40,41]. Even in these well-studied cases, the role of mate choice and signal trait variation within versus between species requires further study, to truly understand the role of sexual selection per se in speciation [42,43].

Most research has focused primarily on the genetic basis of variation in mate choice traits. At the same time, regulatory mechanisms, such as epigenetic modifications, may add an extra layer of information to understand how the genotype is translated into phenotypic variation [44]. However, whether epigenetic modifications are associated with mate choice traits remains poorly understood. This represents a critical gap, as epigenetic variation – particularly DNA methylation – can modulate gene expression, and mediate the interactions between genetic and environmental variation [45,46]. In doing so, epigenetic marks may contribute to phenotypic plasticity and potentially be associated with how mate choice traits vary across environments and express condition-dependent mating signals [25]. Therefore, understanding whether epigenetic variation is associated with mate choice traits might help clarify how genetic and environmental factors interact to shape the evolution of sexual traits involved in divergence and reproductive isolation.

We here focus on DNA methylation, the most extensively studied epigenetic mechanism in ecological and evolutionary research [47,48]. DNA methylation involves the addition of a methyl group to the fifth carbon of a cytosine (5mC) [49]. This molecular modification can affect gene expression, for example, by altering chromatin accessibility and transcriptional activity [50,51]. Notably, DNA methylation can persist through mitotic divisions and, in some cases, through meiotic divisions, potentially transmitting its phenotypic effects to the next generation (e.g., [52–55]).

Variation in DNA methylation is often determined by its genetic background, being genetically programmed to regulate gene activity in some genomic regions [56–59]. At the same time, DNA methylation can also respond to environmental changes, potentially altering expression in response to environmental stimuli – making it an important mediator of phenotypic plasticity [46,60,61]. As a result, DNA methylation and its associated phenotypic effects may arise from genetic effects, environmental effects, or genotype-by-environment interactions (G × E) [45,57].

In this study, we test whether DNA methylation patterns covary with loci associated with variation in mate choice signals. Most molecular work on mate choice has centred on signals rather than mate preference, in part because signals can be easier to measure and more stable than behavioural preferences per se (e.g., [31,35,37]; some notable exceptions on the genetic basis of mate preference aside, e.g. [62]). Thus, investigating the association between variation in DNA methylation and signal traits is a logical place to start to look for a role for epigenetics in sexual isolation. In addition, while several studies have investigated DNA methylation in the context of contributing to sexual dimorphism (e.g., [63–65]), its association with traits involved in mate choice remains largely unexplored.

To test whether DNA methylation is associated with signal traits, we propose a three-step strategy (Fig. 1). First, to genetically map signal traits by identifying loci whose variation is most strongly associated with phenotypic variation. While this approach targets the heritable component of phenotypic variation, the identified genomic regions may also contribute to phenotypic plasticity (e.g., through regulation by DNA methylation). Second, to assess DNA methylation patterns across ecological gradients to identify regions where methylation varies with environmental conditions (*i*.*e*., potentially due to association with locally adapted genetic variants, environmental influences, or genotype-by-environment interactions). Third, to test whether loci associated with signal trait variation are disproportionately enriched in regions exhibiting environmentally-associated DNA methylation. This approach highlights candidate regions where epigenetic marks may contribute to variation in mate choice signals, or otherwise covary with such traits in ways that could affect their evolution.

**Figure 1.**
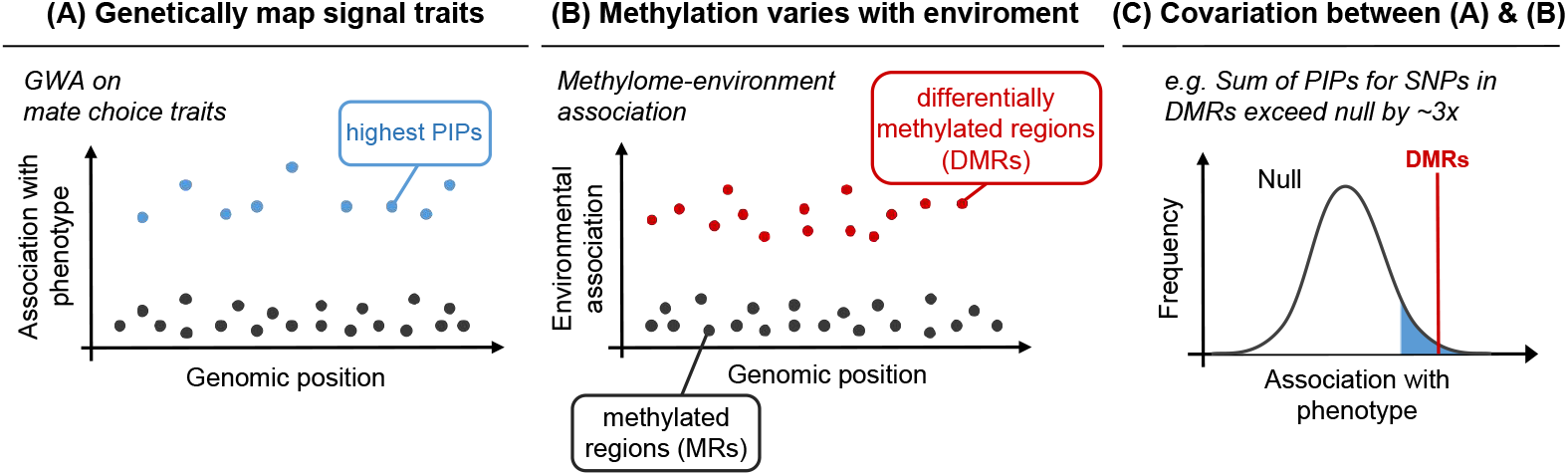
Strategy to test whether environmentally-associated DNA methylation patterns covary with genetic regions associated with mate choice traits. (**A**) Map genetic regions associated with signal traits using genome-wide association studies (GWA). The strength of the association can be measured using statistics such as posterior inclusion probabilities (PIPs) from Bayesian sparse linear mixed models. (**B**) Identify genomic regions for which DNA methylation varies with environmental gradients (i.e., differentially methylated regions, DMRs), for example, using methylome-environment association studies. (**C**) Test for significant associations between loci most strongly associated with signal traits and DMRs. This can be done by evaluating whether DMRs show: (1) higher summed PIPs, (2) higher mean PIPs, and (3) more SNPs in the top 5% of PIPs than expected by chance. *Abbreviation: assoc. = association*.

We apply this framework in *Timema cristinae* stick insects [66]. This species is primarily found on two host-plant species: *Adenostoma fasciculatum* (Rosaceae) and *Ceanothus spinosus* (Rhamnaceae) in chaparral habitats in coastal California, USA (Fig. 2A) [67,68]. Host-associated divergent selection contributes to partial reproductive isolation in this species, giving rise to *Adenostoma* and *Ceanothus* ecotypes [69]. A component of this reproductive isolation is sexual isolation between ecotypes, in which males prefer females from their same ecotype, and prefer conspecific females over heterospecifics [70,71].

**Figure 2.**
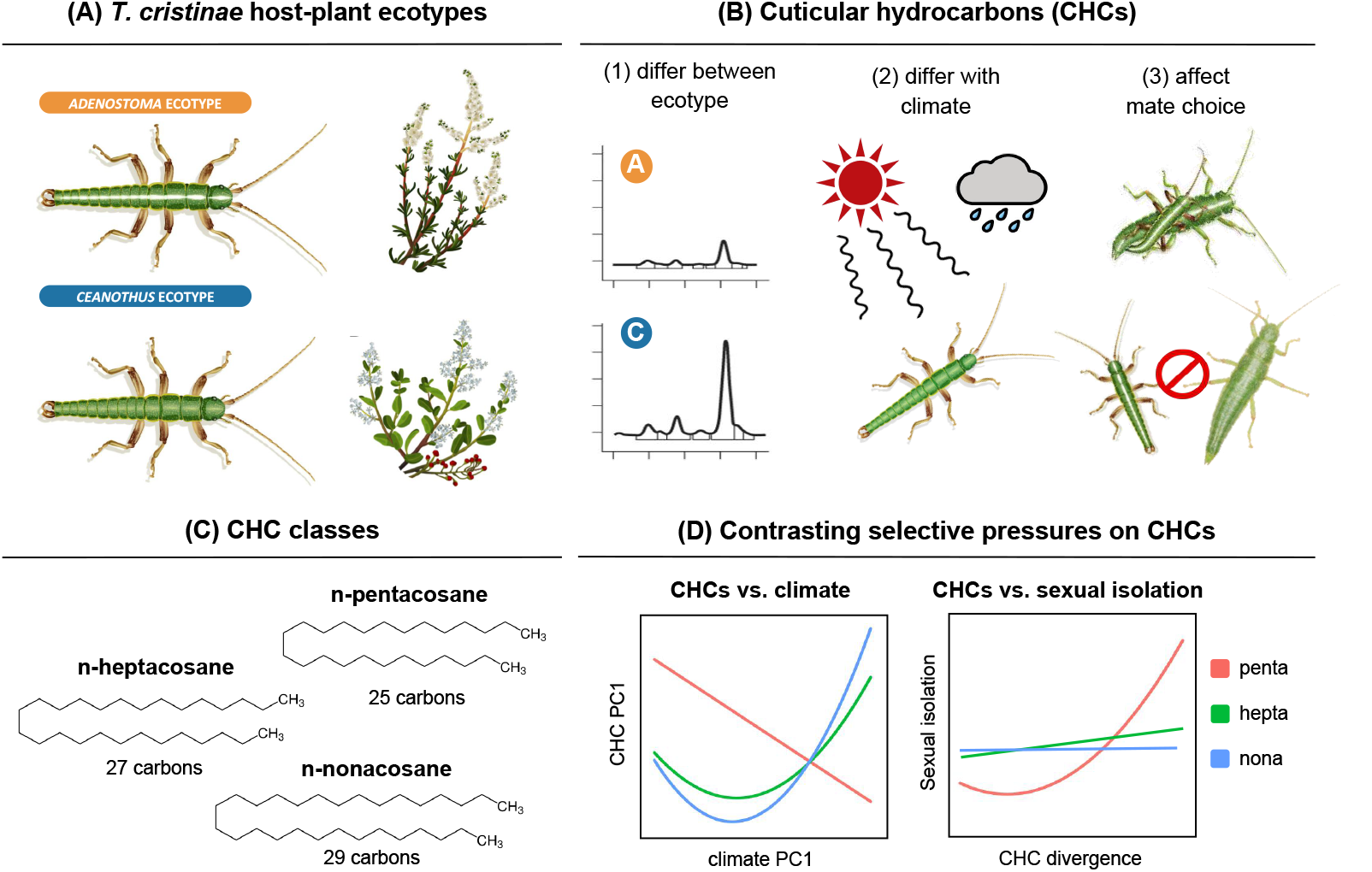
The *Timema cristinae* study system. (**A**) *T. cristinae* and their host plants: *Adenostoma fasciculatum* and *Ceanothus spinosus*. The ecotypes are distinguished by several traits, including colour-pattern, body size, host preference, and cuticular hydrocarbon profiles. (**B**) Properties of cuticular hydrocarbons (CHCs) in *T. cristinae*. CHCs play a key role in desiccation resistance and mate recognition in insects. CHC profiles vary with (1) host-plant ecotype and (2) across climatic conditions; and (3) are used in mate choice and contribute to sexual isolation among *Timema* populations and species. (**C**) Three classes of CHC compounds have been studied in *T. cristinae*, classified by their chain length: penta-, hepta-, and nonacosanes (25, 27 and 29 carbon chains, respectively) **[76-78]**. (**D**) While all CHC classes correlate with climatic variation, divergence in pentacosanes is strongly linked to sexual isolation, whereas hepta- and nonacosanes show no such association **[78]**. The graphs shown were modified from the results of ref. **[78]**.

A key trait underlying both adaptation and sexual isolation in *T. cristinae* is the cuticular hydrocarbons (CHCs) – chemical traits involved in desiccation resistance and mate recognition in insects [72](Fig. 2B). CHCs are often partially plastic, varying with factors like diet and climate [73,74]. Indeed, in *T. cristinae*, these compounds differ between host-plant ecotypes and are associated with climatic differences [75–77]. Although the relative contributions of genetics and environmental induction to these associations remain unclear, CHCs are known to be under polygenic control and exhibit moderate heritability [76,77]. Most importantly, female CHC profiles mediate male mate choice and sexual isolation both within and between species [75]. For example, greater divergence in female CHCs is associated with stronger sexual isolation at both intra- and interspecific levels [75,76]. Moreover, a causal role for CHCs in mate choice has been demonstrated through manipulative perfuming experiments, in which altering CHC profiles affected mate choice within and between species [76]. Furthermore, CHC variation in *T. cristinae* is likely shaped by both natural and sexual selection, with the relative influence of the different forms of selection varying by CHC chain length (Fig. 2C-D). Specifically, shorter-chain compounds (pentacosanes) in *T. cristinae* are most strongly associated with sexual isolation among populations, suggesting that if DNA methylation influences signal traits, the effect might be stronger for pentacosanes than for other CHC traits [78].

Given their polygenic architecture and likely sensitivity to environmental conditions, CHCs are plausible targets for regulation by DNA methylation. To investigate this, we apply the aforementioned approach (Fig. 1) to evaluate whether genomic regions that show differential methylation between ecotypes are enriched for loci associated with CHC variation. Using published genome-wide association studies and methylome-environment association analyses that identified loci associated with CHC variation and ecotype-associated DNA methylation, respectively [76,77,79], we tested for covariation between these genomic regions. By focusing on CHCs as both sexually selected and ecologically relevant traits, our study provides initial insights into the potential mechanistic role of epigenetic modifications in mate choice, and thus of reproductive isolation.

## Material and Methods

### Genomic mapping of variation in CHC traits

We began our investigation focusing on genomic regions associated with CHC variation. To this end, we used results from a published genome-wide association mapping (GWA) study on CHC compounds [77]. In that study, the authors themselves reanalysed previously published genetic and phenotypic data [76] with updated genomic resources.

The phenotypic data used in those studies was obtained as follows. Briefly, Riesch *et al*. [76] collected wild individuals from a single unstructured population (FH, *Adenostoma*, latitude 34.518^°^, longitude -119.801^°^). These individuals were cold-euthanised and had their CHCs extracted by immersing each specimen in 1mL of hexane. A total of 26 CHC compounds were quantified per individual using gas chromatography. The compound abundance was determined by calculating the area under each chromatographic peak, reflecting the relative quantity of each CHC. To allow accurate comparison across samples, a known amount of an internal standard was added to each CHC extract, which was used to scale the areas of the CHC peaks.

Since CHC abundances vary with body size, Riesch *et al*. [76] standardised CHC measurements by calculating the relative abundance of each compound as a proportion of the total CHC profile. The 26 CHC compounds were then categorised into three classes based on their carbon chain length: eight pentacosanes, eight heptacosanes, and ten nonacosanes (*i*.*e*., chains of 25, 27, and 29 hydrocarbons, respectively). The relative proportion of each CHC compound was summed for each CHC class, which was then used as a separate trait for GWA mapping. Genetic variation was assessed using genotype-by-sequencing data for these same phenotyped individuals (175,918 single-nucleotide polymorphisms, SNPs; see ref. [77] for details on mapping to *T. cristinae* reference genome v1.3c2, variant calling, and genotyping).

Following previous work, we focused exclusively on the results for female individuals (n=197), as female CHC profiles have been shown to mediate male mate choice and contribute to sexual reproductive isolation [76,77]. We used results from GWA analyses for each CHC class [77], obtained through Bayesian sparse linear mixed models (BSLMM) implemented in the software *GEMMA* [80]. These models estimate the additive genetic variation of each trait and assign posterior inclusion probabilities (PIP) to individual SNPs. The PIPs represent the probability that a given SNP is associated with phenotypic variation. The GWA results were obtained by running 10 independent Markov chain Monte Carlo (MCMC) chains per trait (1,000,000 sampling steps; 200,000 burn-in; minor allele frequency threshold = 0; see further details on the mapping pipeline in ref. [77]). The proportion of phenotypic variance explained by all SNPs had a median of 89.7% for pentacosanes (95% equal-tail probability interval [ETPI]: 35.8–99.9), 52.5% for heptacosanes (4.9–98.9), and 80.2% for nonacosanes (15.5–99.8) (Table S1).

### *Ecotype-associated differentially methylated regions in* T. cristinae

To investigate ecotype-associated DNA methylation, we used previously published results from ref. [79]. That study assessed DNA methylation differences between host-plant ecotypes of *T. cristinae* using whole-genome bisulfite sequencing. Briefly, the authors obtained whole-methylome data from 24 female individuals (two per population), sampled from 12 populations exhibiting different abundances of *Adenostoma* and *Ceanothus*. Methylation data was mapped to the *T. cristinae* reference genome v1.3c2 [81], and whole-genome sequencing data [82] was used to minimise confounding SNP that could affect methylation calls. Specifically, CpG sites overlapping with C/T and G/A SNPs were removed to improve data accuracy (see ref. [79] for more details).

In the present study, we focused on differentially methylated regions (DMRs) between host-plant ecotypes identified by ref. [79] using methylome-environment association analyses. These analyses applied binomial mixed models (MACAU v1.0.0; [83]) to 1 kilobasepair (kbp) non-overlapping methylation tiling windows (with ≥ 10× coverage per tile; see [79] for more details). In their study, de Carvalho et al. [79] designated DMRs by examining the tail of the empirical *p-value* distribution generated by MACAU, allowing them to assess the properties of DMRs across various cut-off thresholds. Specifically, DMRs were designated based on 10 empirical *p-value* percentiles, varying from the 0.04^th^ to the 0.40^th^ percentile (*P* = 0.0004 to 0.0061), resulting in 25-258 DMRs depending on the cut-off threshold.

We here highlight a few properties of the DMRs designated by ref. [79] that are relevant to this study. First, DMRs showed a stronger association with host-plant ecotype than with genetic, geographic, or climatic variation across the different *p-value* cut-offs. In addition, DMRs were distributed across multiple chromosomes and overlapped at annotated genes at a rate of ∼72%. This proportion resembles the 74% genic overlap observed in the full methylation dataset analysed in MACAU, suggesting that DMRs were not disproportionately localised in genic regions compared to the background distribution of the 1kbp methylation tiles dataset. We used this DMRs dataset to estimate the covariation with genomic regions associated with CHCs output from GWA.

### Identifying associations between host-ecotype DMRs and CHCs’ SNPs

We here tested for associations between DMRs and genomic regions associated with CHC variation. We predicted that DMRs would be enriched for loci associated with each CHC class. We particularly expected the strongest enrichment for loci associated with pentacosanes, due to their connection to mate choice and sexual isolation (Fig. 2D) [78]. To do so, we inferred the degree of association between variation in each female CHC class (pentacosanes, heptacosanes, and nonacosanes) and SNPs using their PIP estimates from the GWA analyses [77]. We then performed two analyses: (1) calculating the total sum of PIP values for SNPs located within DMRs, and (2) calculating the mean PIP values for these SNPs, comparing both metrics against null distributions. Significant increases in either measure relative to null expectations would imply DMRs are enriched for SNPs associated with CHC variation, suggesting a link between DNA methylation and CHC variation (albeit not necessarily a causal link, see Discussion).

We here designated DMRs using four *p-value* cut-offs from the empirical distribution of the methylome-environment association analyses from ref. [79]. We adopted the most stringent (0.04^th^ percentile, *P* < 0.0004), an intermediate (0.20^th^ percentile, *P* < 0.0028), and the more relaxed (0.40^th^ percentile, *P* < 0.0061) thresholds among the *p-value* cut-offs used in that study [79]. We further considered the nominal significance level of *P* < 0.01 as the upper bound of interest. This range of *p-value* cut-offs allowed us to assess the robustness of DMR-CHC associations, serving as a sensitivity analysis.

For each CHC class, we calculated the sum of PIPs for SNPs located within DMRs and compared the observed summed values to a null distribution. To generate this null, we sampled 1,000 random sets of 1kbp methylation tiles (the same as used in methylome-environment association analyses), matched in size and number to the DMRs designated by each *p-value* cut-off. We estimated the empirical *p-value* by calculating the proportion of null sets with the sum of PIPs greater than or equal to the observed values. To quantify effect size, we calculated the observed-to-null PIP ratios (*i*.*e*., the x-fold difference). We applied the same procedure to obtain the mean PIP values to capture average, rather than cumulative, associations between CHC-associated loci and DMRs.

In addition to the sum and mean PIP analyses, we tested whether DMRs were enriched for SNPs exhibiting the top 5% PIPs. This approach evaluates whether loci with the strongest associations to CHC variation are disproportionately located within DMRs compared to random expectations. To do so, we compared the observed number of top 5% PIP SNPs falling within DMRs to a null distribution generated from 1,000 permutations of randomly selected 1kbp methylation tiles, matching the number of DMRs at each *p-value* cut-off. We then calculated empirical *p-values* and x-fold differences. All analyses were conducted both on all DMRs (genic and non-genic) and on genic DMRs only.

To test whether the observed patterns could be explained by differences in minor allele frequencies (MAF) or SNP density within DMRs, we calculated the mean MAF and mean number of SNPs in the DMRs designated at each *p-value* cut-off. These observed values were then compared to null distributions generated using the same permutation approach described above. All statistics were computed separately for each CHC class and *p-value* cutoff. All statistical analyses were performed using R v4.3.2 [84].

## Results

### DMRs are enriched for SNPs associated with CHCs

Consistent with our predictions, SNPs associated with female pentacosanes exhibited significantly higher sum of PIPs within DMRs than expected by chance (∼2 to 7-fold higher, *P* < 0.01 in all comparisons; Fig. 3). In other words, DMRs were enriched for SNPs associated with pentacosane variation. This result was robust across all *p-value* cut-offs (Table 1, Table S2), and remained consistent when considering only genic DMRs (Table S3). Similar results were observed when considering the mean PIPs for SNPs within DMRs (Tables S4-S5). Importantly, we found no significant differences in either the number or the mean MAF of SNPs within DMRs compared to null distributions (Table 1, Table S3), indicating that the observed enrichment was not driven by biases in SNP density or allele frequencies.

**Table 1.**
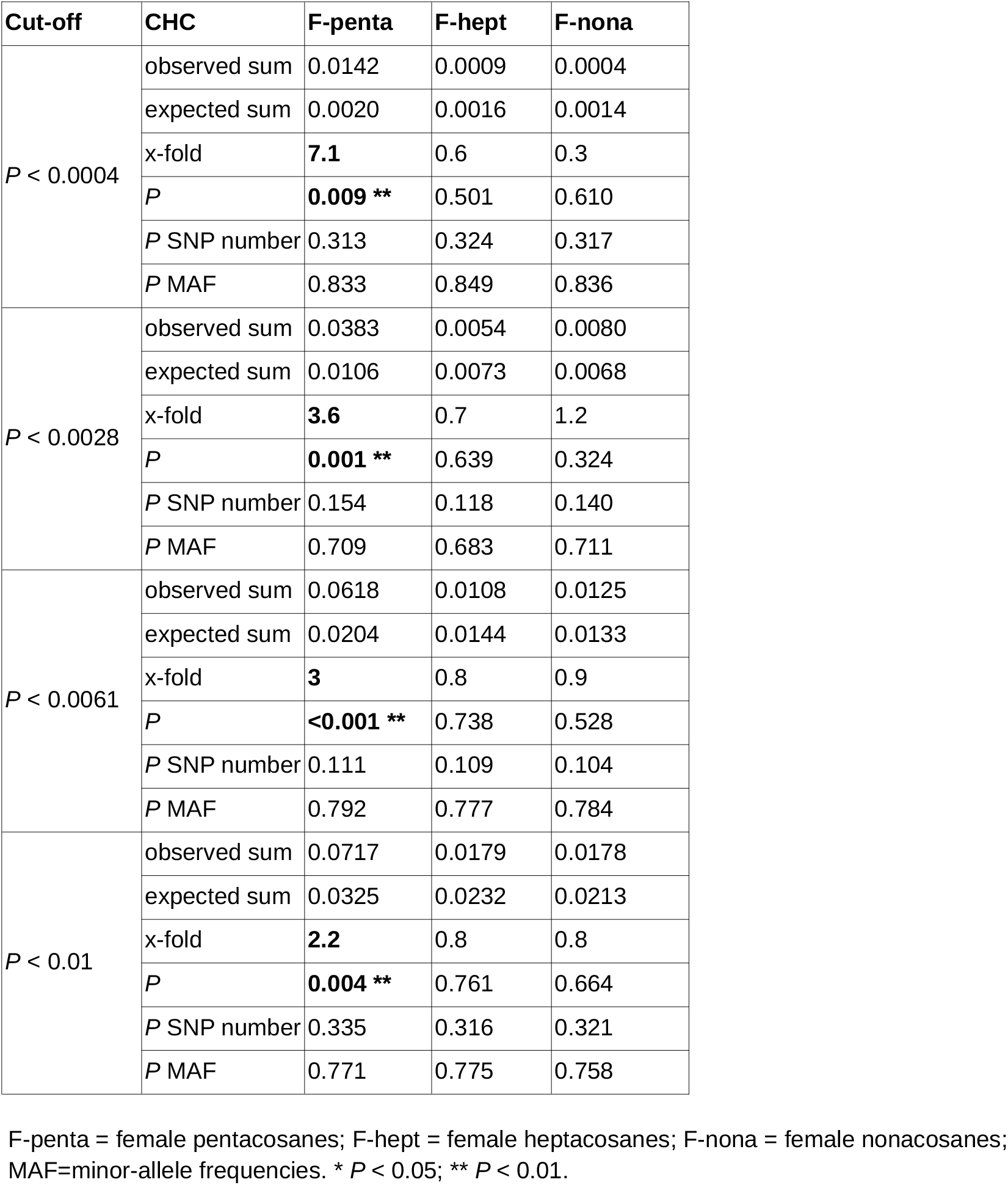
Magnitude of the ratio between the sum of PIPs for SNPs within DMRs compared to null expectations (i.e., x-fold), considering DMRs designated by different p-value cut-offs.

**Figure 3.**
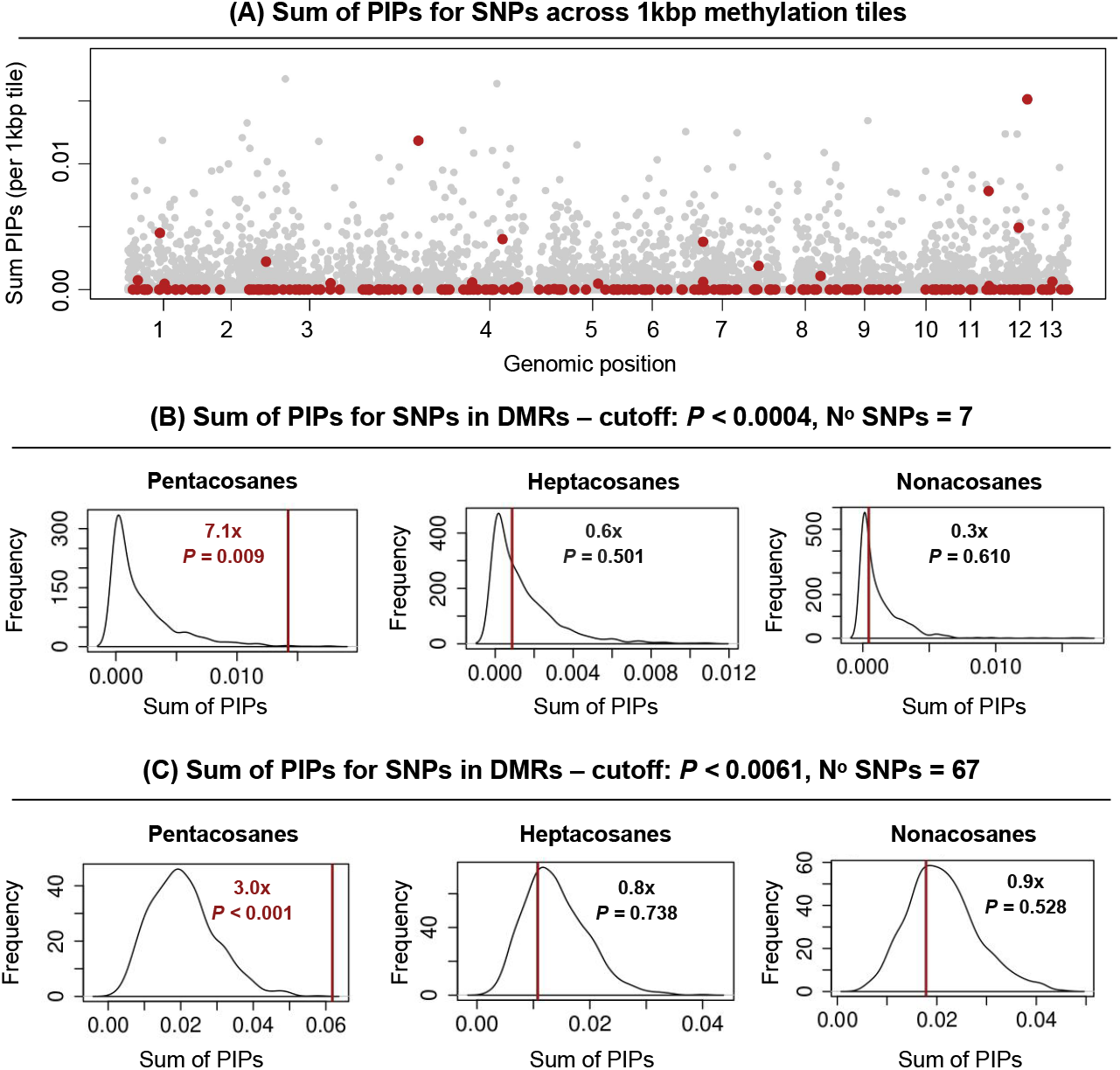
Sum of PIP (posterior inclusion probability) values of SNPs associated with female CHC variation in DMRs compared to genomic background expectations. (**A**) Sum of PIPs across 1 kilobasepair (kbp) methylation tiles, with DMRs highlighted in red. In this graph, DMRs were delimited by the *P <* 0.0061 threshold. The great majority of 1kbp methylation tiles (98%) showed summed PIPs < 0.001. The graphs in (**B**) and (**C**) show the sum of PIPs for SNPs associated with the three CHC classes (i.e., pentacosanes, heptacosanes, and nonacosanes) falling within DMRs (red bars). These observed values are compared to a null distribution generated through randomisations. Each plot shows the results for a different *p-value* cut-off used to delimit DMRs in the methylome-environment association studies (MACAU **[79,83]**), here represented by *P* < 0.0004 and *P* < 0.0061 cut-offs (0.04th and 0.4th percentile of the empirical *p-value* distribution from MACAU; see Table 1). The results consistently show that DMRs are statistically enriched for SNPs linked to female pentacosane variation. Enrichment and *p-values* for these results are shown inside each graph.

Further supporting this association, DMRs defined by more relaxed *p-value* cut-offs (*P* < 0.0061 and *P* < 0.01) showed significant overlap with SNPs in the top 5% of PIPs for pentacosanes (2.7-fold enrichment; *P* < 0.01). This pattern was observed for both all DMRs and those located within genes, whereas more stringent *p-value* cut-offs did not yield significant overlap (0.5–1.5-fold; *P* > 0.10; Tables S6-S7).

In contrast, DMRs showed no significant enrichment for SNPs associated with female heptacosanes or nonacosanes in any analysis, whether considering all DMRs or only genic DMRs (Table 1, Table S3-S7). For example, the sum of PIPs for SNPs associated with hetpacosanes within DMRs was 0.6-0.8-fold lower than expected by chance (*P* > 0.10), and 0.3-1.2-fold in DMRs’ SNPs associated with nonacosanes (*P* > 0.10). Therefore, DMRs tend to show stronger overlap only with genetic regions more significantly associated with female pentacosanes – the class of CHC most strongly implicated in mate choice from past work.

## Discussion

Mate choice arises from preference for signal traits, which could be visual displays, acoustic cues, or chemical signals. Mate choice can reduce gene flow between groups, contribute to reproductive isolation, and help maintain species boundaries [85,86]. As an important agent of sexual selection, it is often expected that mating preference exerts consistent selective pressures on signal traits – though causal direction is often ambiguous and reciprocal evolutionary interactions between traits and preferences are inherent in theories of sexual selection; [16,87]). Although models of sexual selection commonly assume a direct relationship between genotype and phenotype, both preferences and signal trait variation can be partially plastic, varying with environmental conditions [19,22,24]. In this context, investigating epigenetic marks associated with signal traits may help us understand how variation in mating signals arises, and might give us hints about how the environment affects mate choice traits. As such, we here tested whether ecotype-associated DMRs were associated with loci associated with CHC variation, the latter being a known mating signal trait in our study system. Details of our findings follow.

We tested whether DMRs were enriched in SNPs associated with penta-, hepta- and nonacosanes using three metrics: sum, mean or top 5% PIP values of SNPs. All three methods consistently revealed that DMRs are enriched for SNPs associated with pentacosanes. For instance, the sum of PIPs for pentacosane-associated SNPs located within DMRs was 2-to 7-fold higher than expected under the null expectations, indicating significant enrichment. These trends were consistent when considering genic DMRs, suggesting that there might be differential methylation in the genes associated with CHCs. In contrast, we observed no such enrichment for SNPs associated with heptacosanes or nonacosanes. Pentacosanes appear to play an important role in mate choice and sexual isolation in *T. cristinae*. Prior study showed that while divergence in hepta-, and nonacosanes may primarily reflect natural selection from climatic variation (e.g., desiccation resistance and water balance), divergence in pentacosanes is likely shaped by both natural and sexual selection, potentially acting in contrasting directions (Fig. 2D) [78]. Shorter-chained CHCs tend to exhibit lower melting points and higher volatility, likely enhancing the detection of signalling compounds and increasing their prominence in sexual selection, whereas longer-chained ones are typically associated with desiccation resistance [88].

These results thus support our prediction, demonstrating that DMRs were enriched for SNPs specifically associated with the CHC class most strongly linked to attractiveness and mate choice in *T. cristinae*. Since DMRs differ between host-plant ecotypes, DNA methylation may covary with pentacosane-associated genomic regions through three non-mutually exclusive mechanisms. First, genetic determination, where methylation patterns are tied to specific SNPs that affect pentacosane variation (methylation potentially regulates *cis-*genetic variation leading to phenotypic variation) [56,89]. Second, environmental induction, where host-plant habitat directly alters DNA methylation states, which potentially affects pentacosane expression [90,91]. Third, methylation variation reflects a response of genotypes to different host-plant habitats (G x E interactions) [45,92]. In any case, the association between DMRs and pentacosane-associated SNPs suggests that habitat-driven plasticity might contribute to CHC variation.

Our findings align with the idea that mate choice may be influenced by environmental variables and potentially be condition-dependent, where signal traits may convey information about the condition of the sender [20,25]. In this context, DNA methylation could at least serve as a genotype-dependent mechanism that responds to environmental variation, generating different trait expressions that could potentially reflect on the reliability of sexual signalling. This possibility has received very limited attention in the literature. A prior study in black grouse (*Lyrurus tetrix*) found only weak correlations between DNA methylation and melanin-based ornaments (an honest-signalling trait) through sequencing of a single candidate gene [93]. Our study significantly advances this work by showing associations between environmentally-associated DMRs and numerous genomic regions associated with polygenic signal traits in a natural population, revealing a more comprehensive epigenetic architecture associated with sexual signalling. Although CHCs are known to function as condition-dependent sexual signals in other insects such as *Drosophila* [94–96], this remains to be tested more directly in *Timema*. Nevertheless, our findings raise the possibility that sexual signalling traits in this system may be influenced by environmental conditions and mediated by DNA methylation.

Considering that cuticular hydrocarbons are involved in ecotype-specific sexual isolation in *T. cristinae [75,76]*, it is possible that DNA methylation could contribute to this process. However, the extent of this contribution likely depends on the ecological and evolutionary context in which each ecotype occurs. While *T. cristinae* host-plant ecotypes are biologically distinct and differ in multiple traits (including CHC differences), the degree of sexual isolation between them can vary widely [71]. This variation likely reflects a mosaic of factors, including a balance of selection and gene flow (*e*.*g*., driven by geographical distances and climatic differences) and additional traits influencing mate choice beyond CHCs [71,76,97]. Adding to this complexity, previous work shows that DMRs are associated with genetic differentiation among populations, but not specifically with host-plant use [98]. As such, understanding the role of DNA methylation in this complex landscape remains challenging, but may offer key insights into how mate choice traits evolve and contribute to reproductive isolation in nature.

Despite the observed covariation between ecotype-associated DNA methylation patterns and loci associated with pentacosane variation, our results do not establish a causal relationship between methylation and signal trait variation. To build on these findings, future work should directly test whether variation in DNA methylation contributes to differences in pentacosane expression. This could involve experimental or statistical approaches that link DNA methylation levels at DMRs to variation in CHC profiles across individuals with differing genotypes (e.g., through manipulation of DNA methylation and testing for CHC response). In addition, identifying the environmental factors that may trigger methylation changes (e.g., host-plant species) will be key to understanding whether DNA methylation is responsive to environmental change and to what extent it is determined by genetic variation. If a causal relationship between DNA methylation and pentacosane expression is established (whether depending on genetic background or not), the next step would be to evaluate whether methylation-induced changes in CHCs influence mate choice decisions and ultimately reproductive success. Such work would clarify the functional significance of DNA methylation in shaping mating signals, and assess whether methylation contributes to phenotypic plasticity in sexual selection, including condition-dependent signalling. The epigenetic mediation of signal traits further offers the possibility of transmission to the next generation through parental effects [99], which also warrants further study.

Despite ambiguity regarding causal associations, our study is one of the first to establish a link between epigenetic variation and a signal trait of mate choice. This connection offers a promising avenue to understand how environmental variation influences mate choice and how its consequences may shift across space and time. This dynamic view of sexual selection challenges traditional models that assume immutable sexual selection and instead supports a framework where signal expression and mate preferences may be more labile and reflect condition-dependence [19,24,100]. In addition, condition-dependent sexual selection could, in theory, increase the rate and extent of local adaptation and promote population divergence [101]. Our work thus ultimately contributes to a growing understanding of how environmentally responsive sexual selection might influence reproductive isolation and divergence in the wild.

## Acknowledgments

PN and CFdC were supported by Royal Society of London (RG140369). This work was supported by the French Laboratory of Excellence project “TULIP” (ANR-10-LABX-41) funded by a government grant as part of the France 2030 program, as a Senior Package to PN. We gratefully acknowledge the support and computational resources provided by the Center for High Performance Computing at the University of Utah.

