## Supporting Information for "Genomic regions exhibiting divergent methylation patterns covary with loci associated with mate choice traits in a stick insect"

### Supplemental Information

**Table S1. Hyperparameters of genetic mapping of different CHC classes in *T. cristinae.*** In the table, we report proportion of phenotypic variance explained (PVE) by genetic variation, proportion of variance explained by loci with sparse effects (PGE), and the estimated number of SNPs with non-zero (i.e., measurable) effects on the phenotype (n-gamma). Here, we report the median and the 95% equal-tail probability interval in brackets.

| | **PVE** | **PGE** | **n-gamma** |
|---|---|---|---|
| **F-penta** | 0.9 [0.36, 1.00] | 0.16 [0.00, 0.69] | 43 [1, 208] |
| **F-hepta** | 0.53 [0.05, 0.99] | 0.16 [0.00, 0.83] | 14 [0, 230] |
| **F-nona** | 0.8 [0.16, 1.00] | 0.13 [0.00, 0.73] | 22 [0, 163] |

**Table S2. Number of SNPs and DMRs at each *p-value* cut-off used to designate DMRs, considering all DMRs and only genic DMRs.** Numbers in parentheses indicate the number of DMRs that contain at least one SNP.

| **Cut off** | **All DMRs** | | **genic DMRs** | |
|---|---|---|---|---|
| | Nº SNPs | Nº DMRs | Nº SNPs | Nº DMRs |
| ***P* < 0.0004** | 7 | 27 (3) | 7 | 18 (3) |
| ***P* < 0.0028** | 36 | 131 (13) | 29 | 94 (10) |
| ***P* < 0.0061** | 67 | 257 (23) | 54 | 187 (18) |
| ***P* < 0.01** | 88 | 415 (34) | 70 | 299 (26) |

**Table S3. Magnitude of the ratio between the sum of PIPs for SNPs within genic DMRs compared to null expectations (i.e., x-fold), considering DMRs designated by different p-value cut-offs.**

| **cut-off** | **CHC** | **F-penta** | **F-hept** | **F-nona** |
|---|---|---|---|---|
| P < 0.0004 | observed | 0.0142 | 0.0009 | 0.0004 |
| | expected | 0.0014 | 0.0010 | 0.0009 |
| | X-fold | **10.3** | **0.9** | **0.5** |
| | *P* | **0.009 ** ** | 0.5 | 0.61 |
| | P SNP number | 0.16 | 0.18 | 0.16 |
| | P MAF | 0.77 | 0.78 | 0.79 |
| P < 0.0028 | observed | 0.0263 | 0.0039 | 0.0056 |
| | expected | 0.0074 | 0.0051 | 0.0048 |
| | X-fold | **3.6** | **0.8** | **1.2** |
| | *P* | **0.001 ** ** | 0.64 | 0.32 |
| | P SNP number | 0.09 | 0.08 | 0.1 |
| | P MAF | 0.5 | 0.46 | 0.49 |
| P < 0.0061 | observed | 0.0437 | 0.0083 | 0.0086 |
| | expected | 0.0147 | 0.0104 | 0.0095 |
| | X-fold | **3.0** | **0.8** | **0.9** |
| | *P* | **<0.001 ** ** | 0.74 | 0.53 |
| | P SNP number | 0.07 | 0.07 | 0.07 |
| | P MAF | 0.59 | 0.61 | 0.59 |
| P < 0.01 | observed | 0.0537 | 0.0145 | 0.0111 |
| | expected | 0.0237 | 0.0164 | 0.0152 |
| | X-fold | **2.3** | **0.9** | **0.7** |
| | *P* | **0.004 ** ** | 0.76 | 0.66 |
| | P SNP number | 0.2 | 0.19 | 0.2 |
| | P MAF | 0.61 | 0.63 | 0.66 |

F-penta = female pentacosanes; F-hepta = female heptacosanes; F-nona = female nonacosanes; MAF=minor-allele frequencies. * $P < 0.05$; ** $P < 0.01$.

**Table S4. Magnitude of the ratio between the mean PIPs for SNPs within DMRs compared to null expectations (i.e., x-fold), considering DMRs designated by different p-value cut-offs.**

| **Cut-off** | **CHC** | **F-penta** | **F-hepta** | **F-nona** |
|---|---|---|---|---|
| P < 0.0004 | X-fold | **5.2** | 0.44 | 0.25 |
| | *P* | **0.016 *** | 0.672 | 0.765 |
| | P SNP number | 0.317 | 0.345 | 0.308 |
| | P MAF | 0.836 | 0.842 | 0.852 |
| P < 0.0028 | X-fold | **2.6** | 0.52 | 0.85 |
| | *P* | **0.009 **** | 0.878 | 0.557 |
| | P SNP number | 0.142 | 0.136 | 0.152 |
| | P MAF | 0.710 | 0.713 | 0.729 |
| P < 0.0061 | X-fold | **2.28** | 0.55 | 0.7 |
| | *P* | **0.002 **** | 0.951 | 0.827 |
| | P SNP number | 0.091 | 0.095 | 0.093 |
| | P MAF | 0.788 | 0.778 | 0.792 |
| P < 0.01 | X-fold | **2** | 0.72 | 0.76 |
| | *P* | **0.001 **** | 0.895 | 0.831 |
| | P SNP number | 0.339 | 0.324 | 0.324 |
| | P MAF | 0.792 | 0.772 | 0.781 |

F-penta = female pentacosanes; F-hepta = female heptacosanes; F-nona = female nonacosanes; MAF=minor-allele frequencies. * $P < 0.05$; ** $P < 0.01$.

**Table S5. Magnitude of the ratio between the mean PIPs for SNPs within genic DMRs compared to null expectations (i.e., x-fold), considering DMRs designated by different p-value cut-offs.**

| **cut-off** | **CHC** | **F-penta** | **F-hept** | **F-nona** |
|---|---|---|---|---|
| P < 0.0004 | X-fold | **4.95** | 0.43 | 0.23 |
| | *P* | **0.026 *** | 0.612 | 0.712 |
| | *P* SNP number | 0.161 | 0.177 | 0.162 |
| | *P* MAF | 0.771 | 0.78 | 0.793 |
| P < 0.0028 | X-fold | **2.3** | 0.46 | 0.72 |
| | *P* | **0.038 *** | 0.877 | 0.628 |
| | *P* SNP number | 0.087 | 0.082 | 0.099 |
| | *P* MAF | 0.499 | 0.464 | 0.494 |
| P < 0.0061 | X-fold | **2** | 0.55 | 0.61 |
| | *P* | **0.025 *** | 0.91 | 0.85 |
| | *P* SNP number | 0.074 | 0.067 | 0.067 |
| | *P* MAF | 0.594 | 0.613 | 0.589 |
| P < 0.01 | X-fold | **1.82** | 0.74 | 0.59 |
| | *P* | **0.017 *** | 0.82 | 0.928 |
| | *P* SNP number | 0.203 | 0.19 | 0.199 |
| | *P* MAF | 0.613 | 0.626 | 0.659 |

F-penta = female pentacosanes; F-hept = female heptacosanes; F-nona = female nonacosanes; MAF=minor-allele frequencies. * $P < 0.05$; ** $P < 0.01$.

**Table S6: Top 5% PIP values for SNPs within DMRs compared to null expectations.** X-fold indicates the ratio of observed values within DMRs relative to the mean of the null distribution.

| Cuf-off | CHC | penta | hepta | nona |
|---|---|---|---|---|
| P < 0.0004 | observed | 2 | 0 | 0 |
| | mean null | 4.0 | 3.8 | 4.1 |
| | x-fold | 0.5 | 0.0 | 0.0 |
| | *P* | 0.902 | 1.000 | 1.000 |
| P < 0.0028 | observed | 6 | 0 | 2 |
| | mean null | 4.0 | 3.8 | 4.1 |
| | x-fold | 1.5 | 0.0 | 0.48 |
| | *P* | 0.228 | 1.000 | 0.893 |
| P < 0.0061 | observed | 11 | 1 | 3 |
| | mean null | 4.0 | 3.7 | 4.0 |
| | x-fold | 2.7 | 0.3 | 0.8 |
| | *P* | **0.003 *** | 0.967 | 0.728 |
| P < 0.01 | observed | 12 | 3 | 4 |
| | mean null | 4.0 | 3.9 | 4.2 |
| | x-fold | 2.7 | 0.8 | 0.9 |
| | *P* | **0.002 *** | 0.740 | 0.588 |

**Table S7: Top 5% PIP values for SNPs within genic DMRs compared to null expectations.** X-fold indicates the ratio of observed values within genic DMRs relative to the mean of the null distribution.

| **Cuf-off** | **CHC** | **penta** | **hepta** | **nona** |
|---|---|---|---|---|
| P < 0.0004 | observed | 2 | 0 | 0 |
| | mean null | 2.9 | 2.8 | 2.9 |
| | x-fold | 0.7 | 0.0 | 0.0 |
| | *P* | 0.764 | 1.000 | 1.000 |
| P < 0.0028 | observed | 4 | 0 | 1 |
| | mean null | 2..8 | 2.8 | 2.9 |
| | x-fold | 1.4 | 0.0 | 0.4 |
| | *P* | 0.321 | 1.000 | 0.915 |
| P < 0.0061 | observed | 7 | 1 | 2 |
| | mean null | 2.9 | 2.7 | 2.9 |
| | x-fold | 2.4 | 0.4 | 0.7 |
| | *P* | **0.035 *** | 0.914 | 0.768 |
| P < 0.01 | observed | 8 | 3 | 2 |
| | mean null | 2.8 | 2.7 | 2.9 |
| | x-fold | 2.8 | 1.1 | 0.7 |
| | *P* | **0.013 *** | 0.497 | 0.764 |
